# Mitochondrial targeting of glycolysis in a major lineage of eukaryotes

**DOI:** 10.1101/257790

**Authors:** Carolina Río Bártulos, Matthew B. Rogers, Tom A. Williams, Eleni Gentekaki, Henner Brinkmann, Rüdiger Cerff, Marie-Françoise Liaud, Adrian B. Hehl, Nigel R. Yarlett, Ansgar Gruber, Peter G. Kroth, Mark van der Giezen

**Affiliations:** Institute of Genetics, University of Braunschweig, Spielmannstr. 7, D-38106 Braunschweig, Germany.; Biosciences, University of Exeter, Stocker Road, Exeter EX4 4QD, UK.; School of Biological Sciences, University of Bristol, BS81TH, United Kingdom.; Dalhousie University, Department of Biochemistry and Molecular Biology, Halifax, Canada, B3H1X5.; Département de Biochimie, Université de Montréal C.P. 6128, Montréal, Quebec, Canada.; Institute of Parasitology, University of Zürich, Switzerland.; Department of Chemistry and Physical Sciences, Pace University, New York, NY 10038, USA.; Fachbereich Biologie, Universität Konstanz, 78457 Konstanz, Germany.

## Abstract

Glycolysis is a major cytosolic catabolic pathway that provides ATP for many organisms^1^. Mitochondria play an even more important role in the provision of additional cellular ATP for eukaryotes^2^. Here, we show that in many stramenopiles, the C3 part of glycolysis is localised in mitochondria. We discovered genuine mitochondrial targeting signals on the six last enzymes of glycolysis. These targeting signals are recognised and sufficient to import GFP into mitochondria of a heterologous host. Analysis of eukaryotic genomes identified these targeting signals on many glycolytic C3 enzymes in a large group of eukaryotes found in the SAR supergroup^3^, in particular the stramenopiles. Stramenopiles, or heterokonts, are a large group of ecologically important eukaryotes that includes multi- and unicellular algae such as kelp and diatoms, but also economically important oomycete pathogens such as *Phytophthora infestans.* Confocal immunomicroscopy confirmed the mitochondrial location of glycolytic enzymes for the human parasite *Blastocystis.* Enzyme assays on cellular fractions confirmed the presence of the C3 part of glycolysis in *Blastocystis* mitochondria. These activities are sensitive to treatment with proteases and Triton X-100 but not proteases alone. Our work clearly shows that core cellular metabolism is more plastic than previously imagined and suggests new strategies to combat stramenopile pathogens such as the causative agent of late potato blight, *P. infestans.*

---

Mitochondria provide the bulk of cellular ATP for eukaryotes by means of regenerating reduced NAD via the electron transport chain and oxidative phosphorylation^2^. In addition, mitochondria are essential for the production of iron-sulfur clusters^4^, and play roles in heme synthesis, fatty acid and amino acid metabolism^5^. Cytosolic pyruvate is decarboxylated by mitochondrial pyruvate dehydrogenase into acetyl-CoA which enters the citric acid cycle, subsequently producing one GTP (or ATP) and precursors for several anabolic pathways. More importantly, the reduction of NAD^+^ to NADH and production of succinate power the electron transport pathway and oxidative phosphorylation, being responsible for the majority of cellular ATP synthesis. The pyruvate is produced by glycolysis, a widespread cytosolic pathway that converts the six-carbon sugar glucose via a series of ten reactions into the three-carbon sugar pyruvate. Glycolysis is nearly universally present in the cytosol of most eukaryotes but also found in specialised microbodies known as glycosomes originally described in trypanosomatids^6^. More recently, two glycolytic enzymes were also found to be targeted to peroxisomes in fungi due to post-transcriptional processes^7^.

When analysing the genome of the intestinal parasite *Blastocystis^8^*, we discovered putative mitochondrial targeting signals on phosphoglycerate kinase (PGK) and on a fusion protein of triose phosphate isomerase (TPI) and glyceraldehyde phosphate dehydrogenase (GAPDH). The amino-terminal sequences conform to typical mitochondrial targeting signals^9^ and are easily predicted by programmes such as MitoProt^10^. Analysis of the *Blastocystis* TPI-GAPDH and PGK sequences predicts a mitochondrial localisation with high probabilities (P 0.99 and 0.97, respectively). The predicted cleavage sites coincide with the start of the cytosolic enzymes from other organisms (Supplementary Fig. S1A) suggesting that these amino-terminal sequences might target both proteins to the mitochondrial organelle in this parasite^11^. We confirmed the functionality and sufficiency of these putative targeting signals by targeting GFP fused to these signals to mitochondria of a heterologous stramenopile host (Supplementary Fig. S2). Homologous antibodies were raised against *Blastocystis* TPI-GAPDH and PGK to test whether these proteins show an organellar localisation. Compartmentalised distribution of both TPI-GAPDH and PGK was clearly demonstrated using confocal microscopy and 3-dimensional rendering of optical sections (Fig. 1 A-D). Both proteins co-localise with the mitochondrial marker dye MitoTracker and in addition with DAPI which labels the organellar genomic DNA^11,12^ (Fig. 1 E-I).

**Figure 1.**
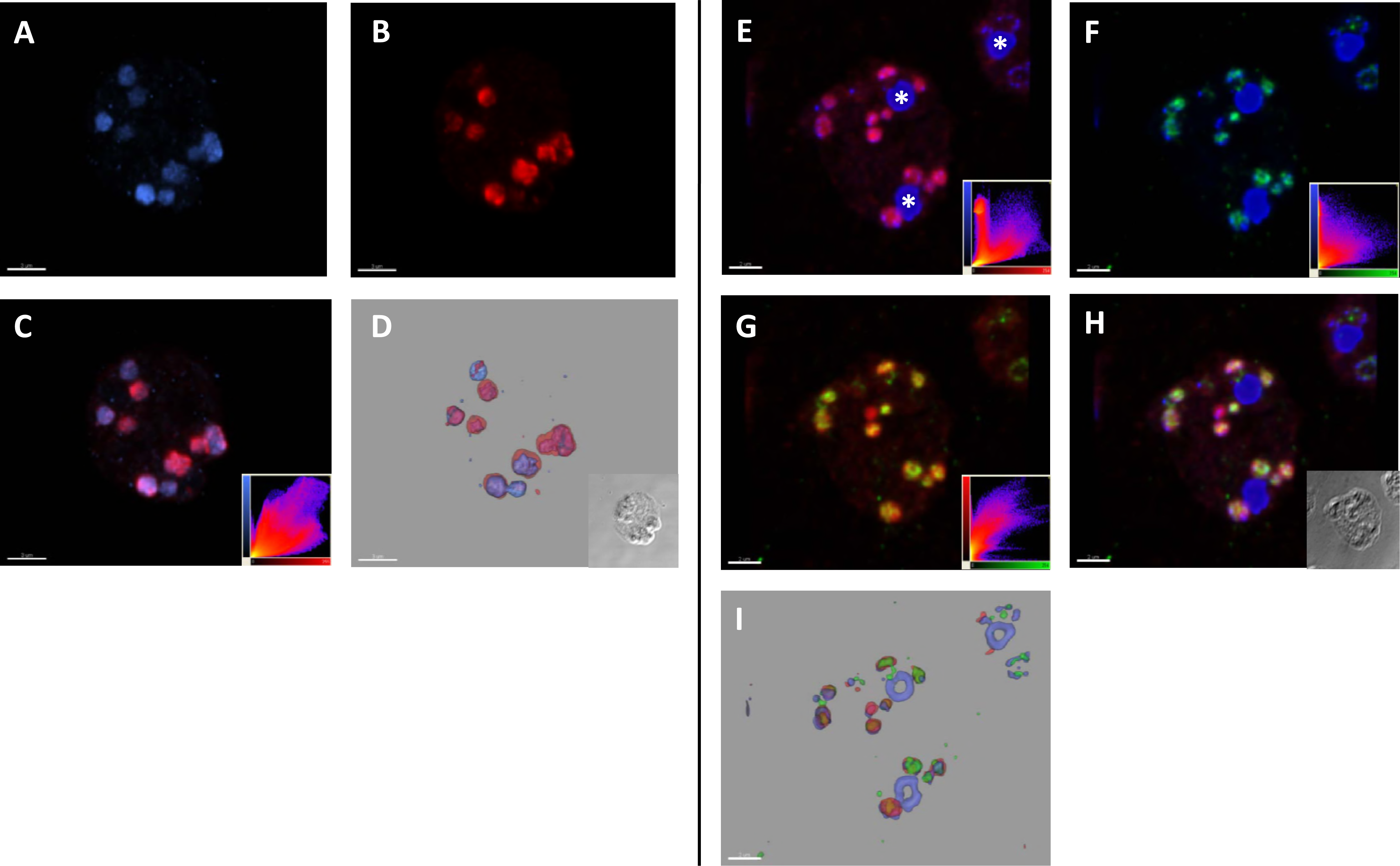
The glycolytic enzymes TPI-GAPDH and PGK localize to mitochondria in the human parasite *Blastocystis.* Three-dimensional immunoconfocal microscopy reconstruction of optical sections (volume rendering) showing representative subcellular localization of PGK (blue) and TPI (red) in trophozoites (A-D). PGK (A) and TPI (B) volume signals show distinct distributions, consistent with localization within mitochondria, with considerable overlap. The merged image (C) provides a qualitative and the scatterplot (inset) of a quantitative measure of signal overlap. Co-localisation of MitoTracker (red) and PGK (green) and DAPI (blue) in trophozoites (E-I). Merged images MitoTracker/DAPI (E) PGK/DAPI (F) and TPI (G) and all three markers together (H) show considerable overlap. Scatterplots (inset) give a quantitative measure of signal overlap for each merged pair of markers (E-G). The DAPI signals (blue) representing nuclear DNA are indicated by asterisks (E). Scale bar 3 μm (A-D) or 2 μm (E-I).

The unexpected mitochondrial localisation of three glycolytic enzymes in *Blastocystis* prompted the analysis of all glycolytic enzymes in this intestinal parasite. Interestingly, targeting signals were only observed on the enzymes of the pay-off phase of glycolysis but not the investment phase (Fig. 2). Although three-dimensional reconstruction of our confocal microscopy data strongly indicated that these enzymes are indeed localised inside *Blastocystis* mitochondria (Fig. 1), we decided to confirm these findings using classical enzyme assays following cellular fractionation. These assays clearly showed that five C3 enzymes are found in the mitochondrial pellet while the five upstream enzymes are all confined to the soluble fraction (Table 1). To assess whether the putative mitochondrial enzymes were only laterally attached to the organelles, as in the case of hexokinase to VDAC in tumours^1^, we tested the latency of enzymatic activities in the presence or absence of Triton X-100. The increase of measurable activity of the C3 enzymes (not shown) suggests they are retained within a membranous compartment. The addition of proteolytic enzymes only affected the measured activity in the presence of the detergent Triton X-100 (Supplementary Table S1), clearly demonstrating that the five C3 glycolytic enzymes in *Blastocystis* are protected by a membrane and reside inside the mitochondria and not on the outside of the organelle, as observed in certain tumours^1^ or as in some proteomics studies^13,14^.

**Figure 2.**
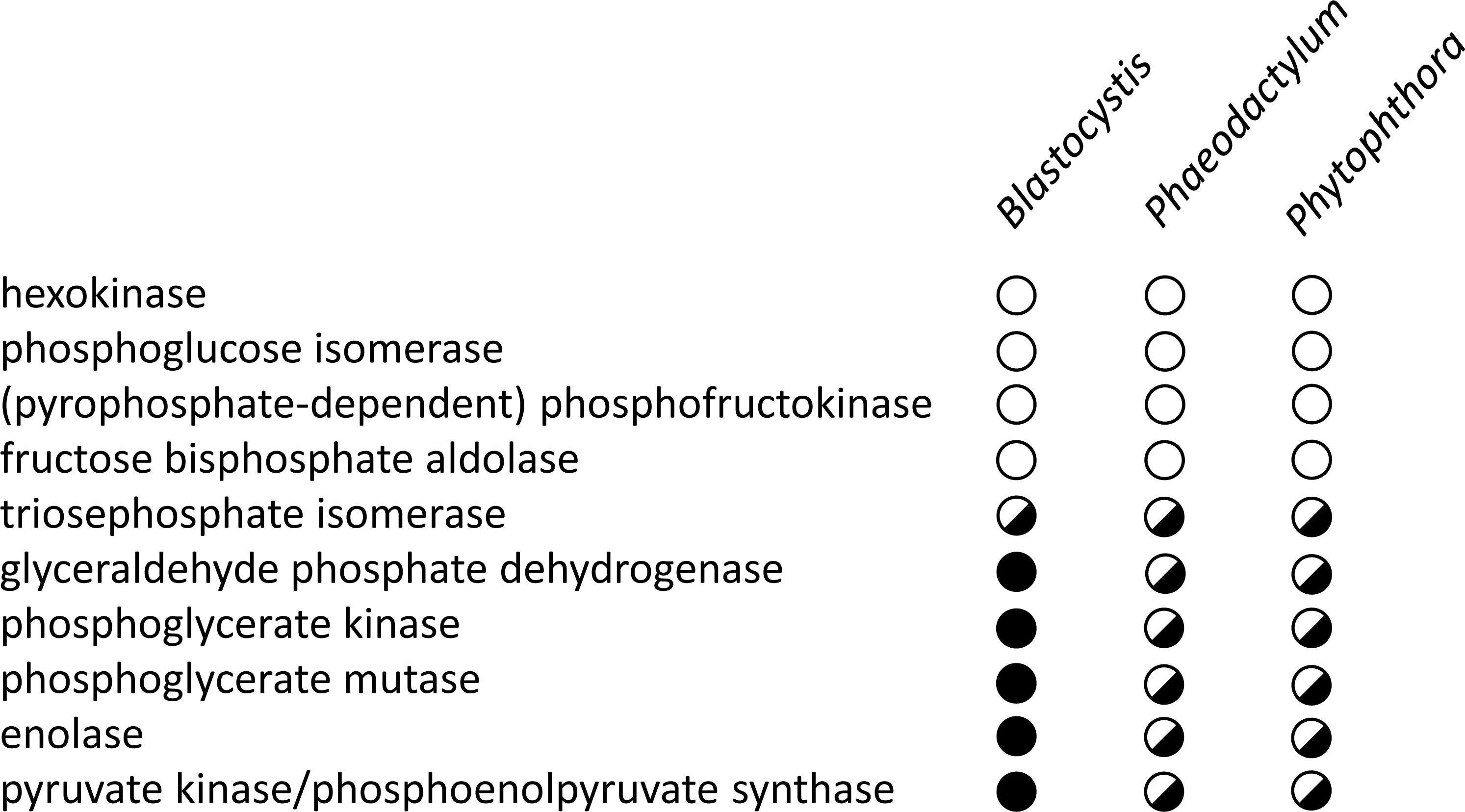
Stramenopile glycolytic enzymes contain mitochondrial-like amino-terminal targeting sequences. Representative stramenopiles with whole genome data known are shown. Presence of mitochondrial-like targeting signal is shown with a filled circle while open circle indicates no mitochondrial-like targeting signal. Where multiple isoforms with and without targeting signal exist, a half-filled circle is shown.

**Table 1.** Pay-off phase glycolytic enzymes in *Blastocystis* are found in the pellet. Activities of glycolytic enzymes from whole cell free extracts (c.f.e.) of *Blastocystis* suspended in phosphate buffered isotonic sucrose solution (pH 7.2). Cells were mixed at a ratio of two volumes of cells: three volumes of 0.5 mm glass beads and broken by three shakes of one minute each at maximum speed on a bead beater (VWR mini bead mill homogenizer (Atlanta, GA, USA)). Cell-free extracts were subjected to increasing centrifugal force producing nuclear, mitochondrial (pellet), lysosomal and cytosolic (supernatant) fractions at 1,912 RCF_av_ for 5 min, 6,723 RCF_av_ for 15 min, 26,892 RCF_av_ for 30 min, respectively. Enzyme activities are the average of three determinations + SD. *\**1 enzyme unit (EU) is the amount of enzyme that converts 1 μmole substrate to product per minute. The yellow box indicates the site of major activity (or in the case of triosephosphate isomerase, the dual localization).

|  | c.f.e. | supernatant | pellet |
| --- | --- | --- | --- |
| hexokinase | 9.3 ± 2.6 | 21.1 ± 3.6 | 2.3 ± 0.6 |
| phosphoglucose isomerase | 3.6 ± 0.8 | 5.8 ± 1.2 | 1.7 ± 0.4 |
| (pyrophosphate-dependent) phosphofructokinase | 11.2 ± 1.8 | 27.1 ± 3.2 | 2.4 ± 1.6 |
| fructose biphosphate aldolase | 3.8 ± 1.3 | 10.9 ± 1.3 | 0.62 ± 0.3 |
| triosephosphate isomerase | 18.3 ± 3.1 | 36 ± 5.1 | 18.0 ± 2.8 |
| glyceraldehyde phosphate dehydrogenase | 9.2 ± 1.4 | 0.5 ± 0.1 | 54.7 ± 5.8 |
| phosphoglycerate kinase | 9.8 ± 1.7 | 5.3 ± 1.3 | 36.2 ± 4.1 |
| phosphoglycerate mutase | 1.2 ± 0.5 | 0.04 ± 0.01 | 2.6 ± 0.7 |
| enolase | 0.42 ± 0.1 | 0.37 ± 0.07 | 1.6 ± 0.4 |
| pyruvate kinase/phosphoenolpyruvate synthase | 2.8 ± 0.7 | 3.3 ± 0.10 | 15.4 ± 2.3 |

As some of us previously reported the mitochondrial localisation of the TPI-GAPDH fusion protein in a related stramenopile^15^, we wondered whether mitochondrial targeting of glycolytic enzymes is more widespread in this group of organisms. When querying all available stramenopile genomes, we noticed the widespread presence of mitochondrial targeting signals on glycolytic enzymes within the whole group. Here, as with *Blastocystis*, only enzymes of the C3 part of glycolysis seem to contain mitochondrial targeting signals (Fig. 2 and supplementary Fig. S1B). To test for functionality, all mitochondrial targeting signals from *Phaeodactylum* C3 glycolytic enzymes were fused to GFP and their cellular location was determined (Supplementary Fig. S2). As with *Blastocystis*, all constructs were targeted to the mitochondrion suggesting these are genuine mitochondrial targeting signals *in vivo*. In addition, we tested mitochondrial targeting signals found on glycolytic enzymes of the oomycete pathogen *Phytophthora infestans*, the water mould *Achlya bisexualis* and the multicellular brown alga *Saccharina latissima*, commonly known as kelp (Supplementary Fig. S2). In all cases, these targeting signals targeted GFP into mitochondria. However, some organisms also contain non-targeting signal bearing glycolytic enzymes suggesting that these cells likely have a branched glycolysis (see Supplementary Fig. S4 for *P. tricornutum).*

Previously, some bioinformatics studies^16,17^ had hinted at the possible mitochondrial location of several glycolytic enzymes. Here, using molecular, biochemical and cell biological methods, we clearly demonstrate the mitochondrial location of glycolytic enzymes of the pay-off phase of glycolysis in a major group of eukaryotes comprising both microbial and multicellular forms. The mitochondrial proteome has a complex and contested evolutionary past^18,19^, and we wondered if glycolytic enzymes targeted to mitochondria might have different evolutionary origins than those that operate in the cytosol. Phylogenetic analysis of all glycolytic enzymes provided no support for this hypothesis because stramenopile glycolytic enzymes cluster with the cytosolic forms of other eukaryotes in phylogenetic trees (Supplementary Figure S3 A-F). This result suggests that the canonical, cytosolic enzymes of glycolysis were targeted to the mitochondrion during stramenopile evolution.

It is difficult to conclusively determine the selective rationale, if any, for the retargeting of glycolysis to stramenopile mitochondria. In *Blastocystis*, and similar to many parasitic eukaryotes^20^, two key glycolytic enzymes have been replaced by pyrophosphate using versions. Normally, the reactions catalysed by phosphofructokinase and pyruvate kinase are virtually irreversible. However, the reactions performed by diphosphate-fructose-6-phosphate 1-phosphotransferase and phosphoenolpyruvate synthase (pyruvate, water dikinase) are reversible, due to the smaller free-energy change in the reaction. As *Blastocystis* is an anaerobe and does not contain normal mitochondrial oxidative phosphorylation^11^, any ATP not invested during glycolysis might be a selective advantage. However, in the absence of these irreversible control points there is a risk of uncontrolled glycolytic oscillations^21^. Separating the investment phase from the pay-off phase by the mitochondrial membrane might therefore prevent futile cycling. However, as not all stramenopiles use pyrophosphate enzymes, this cannot be the whole explanation.

The end-product of glycolysis, pyruvate, is transported into mitochondria via a specific mitochondrial transporter that has only recently been identified^22^ and that is absent from the *Blastocystis* genome^8^. The translocation of the C3 part of glycolysis into mitochondria would necessitate a novel transporter (presumably for triose phosphates). The identification and characterisation of such a transporter would open up new possible drug targets against important pathogens. Examples include *Phytophthora infestans*, the causative agent of late potato blight, which has a devastating effect on food security, but also fish parasites such as *Saprolegnia parasitica* and *Aphanomyces invadans.* Both have serious consequences for aquaculture and the latter causes epizootic ulcerative syndrome, an OIE listed disease^23,24^. Our recent genome analysis of *Blastocystis* identified several putative candidate transporters lacking clear homology to non-stramenopile organisms^8^. Such a unique transporter would not be present in the host (including humans) and could be exploited to prevent, or control, disease outbreaks that currently affect food production while the world population continuous to increase^25^.

## Acknowledgements

The authors wish to thank Professors John F. Allen and Nick Lane (both UCL, UK) for fruitful discussions and criticism. Furthermore, we want to thank Ulrike Brand (TU-BS) and Doris Ballert (Uni KN) for technical assistance. TAW is supported by a Royal Society University Research Fellowship. Work in the lab of MvdG was supported by Wellcome Trust grant 078566/A/05/Z. PGK wishes to acknowledge support by the German Research Foundation (DFG, grant KR 1661/6-1), the Gordon and Betty Moore Foundation GBMF 4966 (grant DiaEdit), and the BioImaging Center of the University of Konstanz as well as Professor Mendel and his group (TU-BS) for using their equipment.

## Materials and methods

### Sources of cDNA and genomic DNA

DNA and cDNA from *Blastocystis* ST1 strain NandII, obtained from a symptomatic human (strain obtained from the American Type Culture Collection, ATCC 50177), was used in this study. Genomic and cDNA libraries of *Phaeodactylum tricornutum* (culture from SAG strain: 1090-1a, Göttingen) were constructed with the "Lambda ZAP II XR library Construction Kit" from Stratagene and the lambda vector EMBL3, respectively. *P. tricornutum* Bohlin (strain 646; University of Texas Culture Collection, Austin) RNA was isolated using TRIzol following manufactures protocol (Thermo Fisher, Germany) and cDNA synthesis was performed with the reverse Transcription system (A3500, Promega, Germany). An *Achlya bisexualis* cDNA library^1^ was kindly provided by D. Bhattacharya (Rutgers University). Screening of libraries, sequencing of positive clones and RACE analyses were performed as described^2^. *Phytophthora infestans* RNA extracted from *P. infestans* mycelia with the RNAeasy Plant Kit from Quiagen and cDNA was synthesized with the Thermo-RT Kit (Display Systems, England). Sequences were also obtained from the EST/genome sequencing programmes from *Phaeodactylum tricornutum^3^* and http://genome.jgi-psf.org/Phatr2/Phatr2.home.html(JGI)^4^, from *Phytophthora infestans* (http://www.pfgd.org^5^) and from *P. sojae* and *P. ramorum* (http://www.jgi.doe.gov^6^).

### GFP constructs for the stable transformation of *Phaeodactylum tricornutum*

Standard cloning procedures were applied^7^. Polymerase chain reaction (PCR) was performed with a Master Cycler Gradient (Eppendorf) using Taq DNA Polymerase (Q BIOgene) according to the manufacturer’s instructions. cDNA from *Blastocystis* ST1 strain NandII (Bl), *Phaeodactylum tricornutum* (Pt), *Phytophthora infestans* (Pi) and *Achlya bisexualis* (Ab) was used as template for the PCR reactions. For *Saccharina latissima* (Sl) a cDNA clone (ABU96661) was used as template.

PCR products were cloned into TA-vector PCR 2.1 (Invitrogen) or blunt cloned into pBluescript II SK^+^ (Stratagene). The primers (Table 1) allowed insertion of restriction enzyme recognition sites (*Eco*RI/N*coI* or *Sma*I/N*coI*) that were used to clone the presequences in frame to eGFP within pBluescript-GFP. The presequence-GFP fusions were cut out with appropriate restrictions enzymes *(EcoRI/HindIII* or SmaI/HindIII) and cloned into the *Phaeodactylum tricornutum* transformation vector pPha-T1^8,9^, either into the corresponding sites or, in case of SmaI, into the EcoRV site. For the constructs with Protein ID (Fig. S4) a slightly different cloning approach was used. PCR with a proof reading Polymerase (*Pfu* or Kapa Hifi) were used to amplify corresponding fragments from cDNA. These fragments were cloned blunt end in a modified pPha-T1 Vector. These Vectors include an eGFP with a StuI or *KspAI* restriction site, allowing a one-step cloning procedure, with subsequent screening for the correct orientation of the fragment at the N-terminus of eGFP. The *Blastocystis* presequences were produced by kinasing the primers using T4 polynucleotide kinase using manufacturer’s procedures and subsequently annealing in a thermal cycler after which they were cloned into the diatom expression vector equipped with eGFP and the StuI restriction site.

**Table 1:**
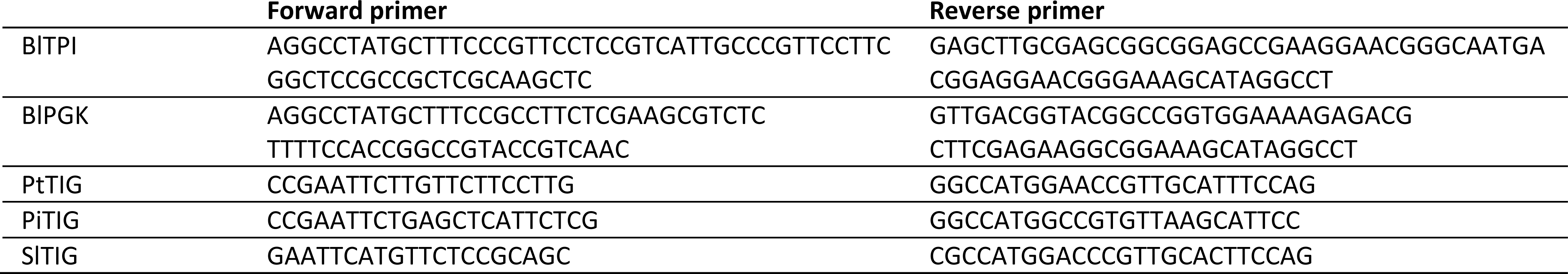
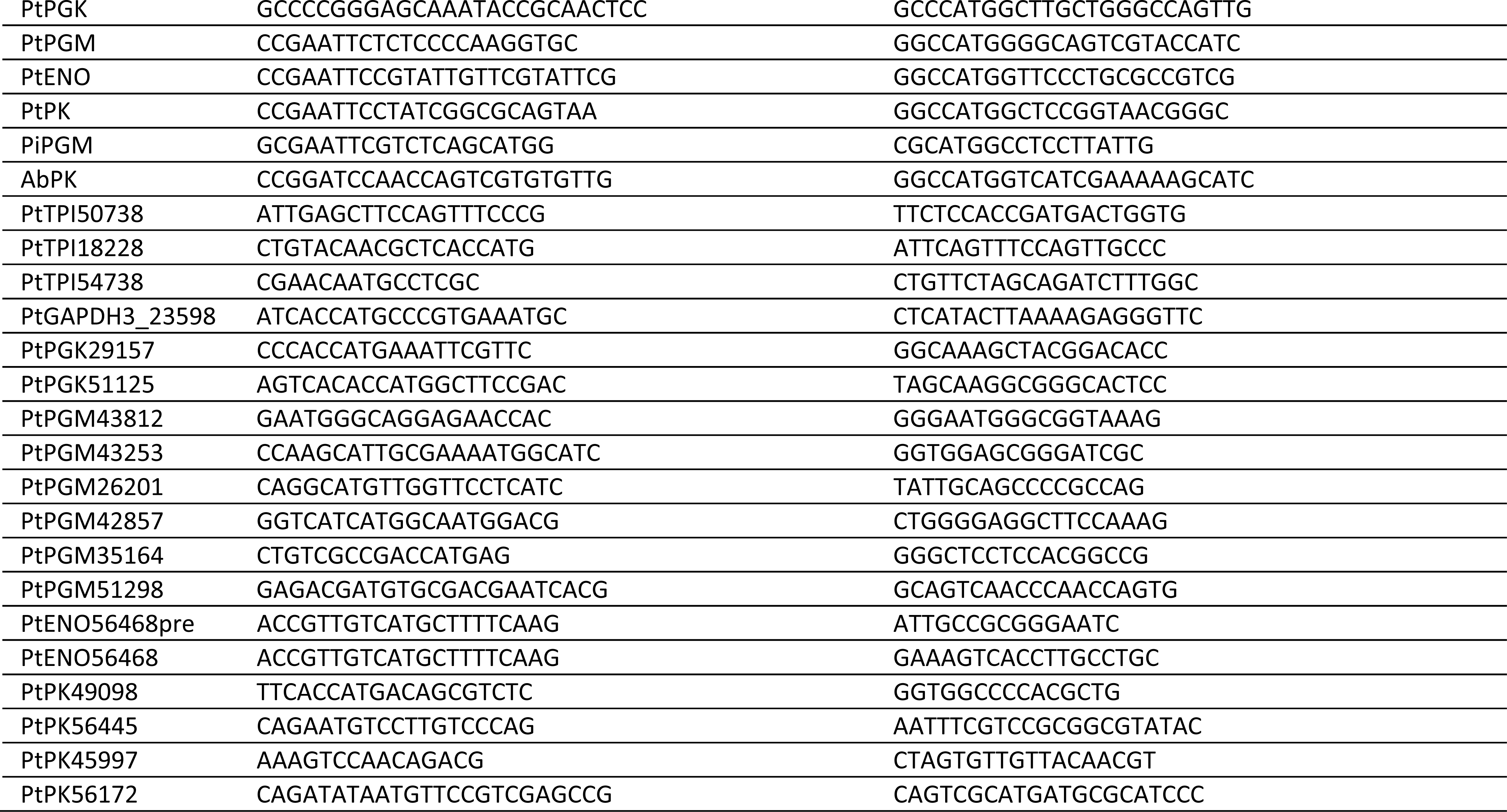
PCR primers for production of GFP fusion constructs.

### Transformation of *Phaeodactylum tricornutum*

*Phaeodactylum tricornutum* Bohlin (UTEX, strain 646) was grown at 22 °C under continuously light of 75 μE in artificial seawater (Tropic Marin) at a Q.5 concentration. Transformations were performed as described by Zaslavskaia *et al*.^8,10^ For each transformation, tungsten particles. For each transformation, tungsten particles M1Q (Q.7 μm median diameter) covered with 7-20 μg DNA were used to bombard cells with the Particle Delivery System PDS-1000 (Bio-Rad, HE-System) prepared with 650, 900, 1100 or 1350 psi rupture discs.

### Microscopic analysis of transformed *Phaeodactylum tricornutum*

Reporter gene expression was visualized using confocal laser scanning microscopy (cLSM-510META, Carl Zeiss, Jena, Germany) using a Plan-Neofluar 40x/1.3 Oil DIC objective. The eGFP fusion proteins were excited with an argon laser at 488 nm with 8-10% of laser capacity. Excited fluorophores were detected with a bandpass filter GFP (505-530 nm) using a photomultiplier. Chlorophyll *a* autofluorescence was simultaneously detected with a META-channel (644-719 nm). MitoTraker Orange CM-H_2_TMRos (Molecular Probes) was applied for fluorescence staining of mitochondria. *P. tricornutum* cells were stained according to the protocol of the manufacturer. Cells were incubated with 100 nM dye solution, incubated for 30 minutes, washed and observed (images were recorded using the Multitracking mode with the following parameters for Wavelength T1 = 488 nm 10% and T2 = 543 nm 100% laser line, primary beam splitting mirrors UV/488/543/633 nm; emitted light was detected with the META-channel).

### Protein production and antibody generation

*Blastocystis* TPI-GAPDH was amplified from cDNA using primers TPI-GAPDH pET F: aga aga *CAT ATG* TTC GTC GGT GGC AAT TGG AAG TGC AA and TPI-GAPDH pET R: tct tct *GGA TCC* TTA AGA GCG ATC CAC CTT CGC CA adding *NdeI* and BamHI restriction sites, respectively, to facilitate cloning in gene expression vector pET14b (Novagen, Merck, Whatford, UK). The *Blastocystis* PGK was amplified from cDNA using PGK pET F: aga aga *CAT ATG* AAG CTG GGA GTT GCT GCC TAC G and PGK pET R: tct tct *CAT ATG* TCA CGC GTC CGT CAG AGC GGC CAC ACC C which added NdeI restriction sites for pET14b cloning. The mitochondrial targeting signals were not amplified as these would not be part of the mature processed protein. All constructs were confirmed by sequencing. The in-frame His-tag allowed for affinity chromatography purification of the recombinant protein. Recombinant *Blastocystis* TPI-GAPDH and PGK were used to immunise guinea pigs and rabbits, respectively, for polyclonal antibody generation at Eurogentec (Seraing, Belgium).

### Culture conditions for *Blastocystis*

*Blastocystis* isolate B (originally designated *Blastocystis* sp. group VII^11^, now called ST7^12^) was used. The parasite was grown in 10 ml pre-reduced Iscove’s modified Dulbecco’s medium (IMDM) supplemented with 10% heat-inactivated horse serum. Cultures were incubated for 48 h in anaerobic jars using an Oxoid AneroGen pack at 37 ^o^C. Two-day-old cultures were centrifuged at 1600 g for 10 min, washed once in a buffer consisting of 30 mM potassium phosphate, 74 mM NaCl, 0.6 mM CaCl_2_ and 1.6 mM KCl, pH 7.4 and resuspended in an a nitrogen gassed isotonic buffer consisting of 200 mM sucrose (pH 7.2) containing 30 mM phosphate, 15 mM mercaptoethanol, 30 mM NaCl, 0.6 mM CaCl_2_, and 0.6 mM KCl (pH 7.2).

### Subcellular fractionation of *Blastocystis*

*Blastocystis* cells were broken by mixing 2 volumes of the cell suspension with 3 volumes of 0.5 mm beads and broken by 3 one minute duration shakes at maximum speed on a bead breaker (VWR mini bead mill homogenizer, Atlanta, GA, USA) with one-minute pauses on ice. Cell-free extracts were subjected to increasing centrifugal force producing nuclear (N, 1,912 RCF_av_ for 5 min), mitochondrialike (ML, 6,723 RCF_av_ for 15 min), lysosomal (L, 26,892 RCF_av_ for 30 min) and cytosolic (S) fractions, respectively, using a using a Sorvall RC-2B centrifuge fitted with an SS-34 rotor.

### Enzyme assays

Hexokinase was assayed by measuring the reduction of NAD^+^ at 340 nm in a coupled reaction with *Leuconostoc mesenteroides* glucose-6-phophate dehydrogenase (3 EU), containing 38 mM Tris-HCl pH 7.6, 115 mM D-glucose, 10 mM MgCl_2_, 0.5 mM ATP, 0.2 mM NAD^+^, 0.05 mL of *Blastocystis* cell-free extract (0.08-0.12 mg) or fraction (N, 0.15-0.18 mg; ML, 0.12-0.17 mg; L, 0.08-0.11 mg; S, 0.090.05 mg), in a final volume of 1 mL at 25 ^o^C.

Phosphoglucose isomerase was assayed by measuring contained g the reduction of NADP^+^ at 340 nm in a coupled reaction with *Leuconostoc mesenteroides* glucose-6-phophate dehydrogenase (2 EU), containing 38 mM Tris-HCl pH 7.6, 3.3 mM D-fructose-6-phosphate, 0.66 mM ß-NADP^+^, 3.3 mM MgCl_2_, 0.05 mL of *Blastocystis* cell-free extract or fraction in a final volume of 3 mL at 25 ^o^C.

Phosphofructokinase was assayed using the standard coupled assay containing 38 mM Tris-HCl pH 7.6, 5 mM dithiothreitol, 5 mM MgCl_2_, 0.28 mM NADH, 0.1 mM ATP, 0.1 mM AMP, 0.8 mM fructose-6-ohosphate, 0.4 mM (NH_4_)_2_SO_4_, 0.05 EU each of rabbit muscle aldolase, rabbit muscle glycerophosphate dehydrogenase, and rabbit muscle triosephosphate isomerase, 0.05 mL of *Blastocystis* cell-free extract or fraction in a final volume of 3 mL at 25 ^o^C.

Aldolase was assayed using a modification of the hydrazine method in which 3-phosphoglyceraldehyde reacts with hydrazine to form a hydrazone which absorbs at 240 nm; the assay contained 12 mM fructose-1,6-bisphosphate, pH 7.6, 0.1 mM EDTA, 3.5 mM hydrazine sulfate and 0.05 mL of *Blastocystis* cell-free extract or fraction in a final volume of 3 mL at 25 ^o^C.

Triosephosphate isomerase was assayed by measuring the oxidation of NADH using a linked reaction with glycerol-3-phosphate dehydrogenase; 220 mM triethanolamine pH 7.6, 0.20 mM DL-glyceraldehyde-3-phosphate, 0.27 mM NADH, 1.7 EU glycerol-3-phosphate dehydrogenase, and 0.05 mL of *Blastocystis* cell-free extract or fraction in a final volume of 3 mL at 25 ^o^C.

Glyceraldehyde-3-phosphate dehydrogenase was assayed by measuring the initial reduction of NAD^+^ at 340 nm; the assay contained 13 mM sodium pyrophosphate pH 8.0, 26 mM sodium arsenate, 0.25 mM NAD, 3.3 mM dithiothreitol, and 0.05 mL of *Blastocystis* cell-free extract or fraction in a final volume of 3 mL at 25 ^o^C.

Phosphoglycerate kinase was assayed by measuring the 3-phosphoglycerate dependent oxidation of NADH at 340 nm; the assay contained 40 mM Tris-HCl pH 8.0, 0.5 mM MgCl_2_, 0.26 mM NADH, 0.1 mM ATP, 2 EU *S. cerevisiae* glyceraldehydephosphate dehydrogenase, and 0.05 mL of *B. hominis* cell free extract or fraction in a final volume of 3 mL at 25 ^o^C.

Phosphoglycerate mutase was measured using the standard coupled assay and measuring the decrease in absorbance at 340 nm; the assay contained 76 mM triethanolamine pH 8.0, 7 mM D(-) 3-phosphoglyceric acid, 0.7 mM ADP, 1.4 mM 2,3-diphosphoglyceric acid, 0.16 mM NADH, 2.6 mM MgSO_4_, 100 mM KCl, 5 EU pyruvate kinase/8 EU lactate dehydrogenase from rabbit muscle, 5 EU rabbit muscle enolase, and 0.05 mL of *Blastocystis* cell-free extract or fraction in a final volume of 3 mL at 25 ^o^C.

Enolase was determined using the standard coupled assay and measuring the decrease in absorbance at 340 nm; the assay contained 80 mM triethanolamine pH 8.0, 1.8 mM D(+) 2-phospholycerate, 0.1 mM NADH, 25 mM MgSO_4_, 100 mM KCl, 1.3 mM ADP, 5 EU pyruvate kinase/8 EU lactate dehydrogenase from rabbit muscle, and 0.05 mL of *Blastocystis* cell-free extract or fraction in a final volume of 3 mL at 25°C.

Pyruvate kinase was determined by measuring the oxidation of NADH at 340 nm using the following mixture, 45 mM imidazole-HCl pH 8.0, 1.5 mM ADP, 0.2 mM NADH, 1.5 mM phosphoenolpyruvate, 5 EU rabbit muscle lactate dehydrogenase, and 0.05 mL of *Blastocystis* cell-free extract or fraction in a final volume of 3 mL at 25 ^o^C.

Pyruvate phosphate dikinase was assayed spectrophotometrically by measuring the oxidation of NADH at 340 nm in 3 mL cuvettes. The reaction contained HEPES buffer (pH 8.0), 6 mM MgSO_4_, 25 mM NH_4_Cl, 5 mM dithiothreitol, 0.1 mM disodium pyrophosphate, 0.25 mM AMP, 0.1 mM phosphoenolpyruvate, and 0.05-0.25 mg of *Blastocystis* cell-free extract or fraction. The rate of pyruvate production was determined by the addition of 2 U of lactate dehydrogenase and 0.25 mM NADH, and compared to controls with phosphoenolpyruvate but lacking AMP, and those containing AMP but lacking phosphoenolpyruvate. The concentration of AMP, pyrophosphate and phosphoenolpyruvate used in the assay was selected from preliminary assays using varying concentrations from 0.025-1.0 mM. The generation of ATP from AMP by pyruvate phosphate dikinase was confirmed by measuring the ATP formed using a luciferin/luciferse assay (Molecular Probes, In Vitrogen, Eugene, OR, USA). The assay was performed as described above but lacking lactate dehydrogenase and NADH, after varying times 0, 15, 30, 45 and 60 min 0.1 mL of the assay is removed and added to one well of a 96 well plate containing 0.1 mL of 0.25 |j,g firefly luciferase and 0.5 mM luciferin and the luminescence recorded using a Spectra Max M2 plate reader (Molecular Devices, Sunnyvale, CA).

The activity of pyrophosphate dependent phosphofructokinase* in the direction of fructose-1,6-bisphosphate formation (forward reaction) was determined in 1 mL assay volumes containing 0.1 M HEPES-HCl, pH 7.8; 20 mM fructose-6-phosphate; 2 mM Na pyrophosphate; 5 mM MgCl_2_; 0.25 mM NADH; 0.2 U of aldolase (from rabbit muscle); and 0.3 U each of glycerophosphate dehydrogenase (from rabbit muscle) and triosephosphate isomerase (from rabbit muscle), 10 |J,M fructose 2,6 diphosphate. The reaction was initiated by addition of 0.05-0.25 mg of *Blastocystis* cell-free extract or fraction, and the rate of NADH oxidation was followed at 340 nm on a Beckman DU 640 spectrophotometer (Indianapolis, IN, USA). The activity of the reverse reaction was determined by measuring orthophosphate-dependent formation of fructose-6-phosphate from fructose-1,6-bisphosphate. The reaction mixture (1 mL) contained 0.1 M HEPES-HCl, pH 7.8; 2 mM fructose-1,6-bisphosphate; 15 mM NaH2PO4; 5 mM MgCl2; 0.3 mM NADP^+^ and 0.12 U glucose-6-phosphate dehydrogenase and 0.24 U glucose phosphate isomerase. The reaction was initiated by addition of 1 mg of pyrophosphate dependent phosphofructokinase and monitored at 340 nm. *Pyrophosphate fructose-6-phosphate 1-phosphotransferase (PFP).

### Confocal microscopy of *Blastocystis*

*Blastocystis* trophozoites were treated with MitoTracker Red (Molecular Probes), washed, fixed in 10% formalin and incubated in ice cold acetone for 15 minutes and air-dried.

Slides with fixed parasites were rehydrated in phosphate buffered saline (PBS) for 30 minutes and blocked with 2% BSA in PBS for 1 hour at room temperature. All antibody incubations were performed at room temperature in 2% BSA in PBS, 0.1% triton X-100. Slides were washed 5 times in 0.2 % BSA in PBS, 0.01% triton X-100 between incubations to remove unbound antibodies.

Primary antibodies: Rabbit, anti-PGK; Guinea Pig, anti-TPI-GAPDH (Eurogentec, Seraing, Belgium) were used at a dilution of 1:500 and 1:300 in 2% BSA in PBS, 0.1% triton X-100, respectively.

Secondary antibodies: Alexa Fluor 488 conjugated Goat anti-Rabbit (Invitrogen, Eugene, OR, USA), Alexa Fluor 405 conjugated Goat anti-Rabbit (Invitrogen, Eugene, OR, USA), TRITC-conjugated Goat anti-Guinea Pig were used at 1:200 dilutions in 2% BSA in PBS, 0.1% triton X-100, each.

The DNA intercalating agent 4’-6-Diamidino-2-phenylindole (DAPI) for detection of nuclear and mitochondrial DNA was added to the final but one washing solution at a concentration of 1 μg-ml^−1^. The labeled samples were embedded in Dako Glycergel Mounting Medium (DAKO, Carpinteira, CA, USA) and stored at 4 °C.

Immunofluorescence analysis and image data collection was performed on a Leica SP2 AOBS confocal laser-scanning microscope (Leica Microsystems, Wetzlar, Germany) using a glycerol immersion objective lens (Leica, HCX PL APO CS 63x 1.3 Corr). Image z-stacks were collected with a pinhole setting of Airy 1 and twofold oversampling. Image stacks of optical sections were further processed using the Huygens deconvolution software package version 2.7 (Scientific Volume Imaging, Hilversum, NL). Three-dimensional reconstruction, volume and surface rendering, and quantification of signal overlap in the 3D volume model were generated with the Imaris software suite (Version 7.2.1, Bitplane, Zurich, Switzerland). The degree of signal overlap in the 3D volume model is depicted graphically as scatterplots. The intensity of two fluorescent signals in each voxel of the 3D model is measured and plotted. Voxels with similar signal intensity for both signals appear in the area of the diagonal. All image stacks were corrected for spectral shift before rendering and signal colocalization analysis.

### Phylogenetic analyses

Sequences of all glycolytic enzymes from *Phaeodactylum tricornutum* and *Blastocystis* ST1, strain NandII, were used as seeds in BlastP searches in the non-redundant database at the NCBI^13^. We were especially interested to identify all sequences in the SAR supergroup^14^ (Stramenopiles, Alveolates and Rhizaria). In addition, representatives from other eukaryotic groups were added and, if required, closely related bacterial sequences. Sequences were automatically added to pre-existing alignments and subsequently manually refined using the Edit option of the MUST package^15^. Final datasets were generated after elimination of highly variable regions and positions with more than 50% gaps by G-blocks^16^. All datasets were first analysed with a maximum likelihood (ML) method under two different models. PhyML v2.3^17^ was used with the SPR moves option and the LG+F+4G model^18^ and PhyML v3 (with SPR moves) was used using the C20+4G model, corresponding to 20 pre-calculated fixed profiles of positional amino-acid substitution^18^. Based on the likelihood values (l), the number of parameters (K) and alignment positions (n), the AIC (AIC= -2l +2K) and the corrected AIC (AICc; AIC+ 2K(K+1)/n-K-1) was calculated^19^. The lowest AICc value corresponds to the best tree, if the value of the C20 analysis was better, then a second ML analysis under the C40+4G model was performed and the AICc value estimated, until the overall best model was found. If the AICc of C40 is better than C20 then C60 was tested. Once the best model was estimated for all six datasets, a rapid bootstrap analysis with 100 replicates in RAxML v7 under the LG model was performed^20^ and an additional analysis in Phylobayes v3 with the CATfix C20 model in all cases or, alternatively, the best C-model. Two independent chains were run for 10,000 points and trees are sampled at every tenth points^21^. Trees obtained with the best model are presented and both posterior probabilities (PP) and rapid bootstrap values (BS) are indicated on trees if PP>0.5 or BS >30%, respectively.

### Cellular localisation predictions

TargetP^22^ and MitoProt^23^ were used to analyse putative subcellular localization. Using the non-plant and no cut-offs settings. In case of Viridiplantae, Rhodophyta and Glaucocystophyta the plant results were taken, if non-plant results differ.

## Amino acid sequences of mitochondrial targeting sequences used in GFP targeting experiments as seen in Supplementary Figure S2

**A.**

~~~
>preTPI-GAPDH-GFP *(Blastocystis)* (OAO12326)
MLSRSSVIARSFGSAARKL
>prePGK-GFP *(Blastocystis)* (OAO15536)
MLSAFSKRLFSTGRTVN
~~~

**B.**

~~~
>preTPI-GAPDH-GFP *(Phaeodactylum)* (NCBI AF063804)
MLASSRTAAASVQRMSSRAFHASSLTEARKFFVGGNWKCNGS
>prePGK-GFP *(Phaeodactylum)(JGI* 48983)
MLFRMLTSTALRRSPVTTSLTCCCKANAFAVRIRSFHAAPVIQAKMTVEQLAQQ
>prePGM-GFP *(Phaeodactylum)(JGI* 33839)
MFAVSRSSFLLATRVKTLRSFAAVQAADKHTLVLLRHGESTWNLENKFTGWYDCP
>preENO-GFP *(Phaeodactylum)(JGI* 1572)
MMWSRPVLRRNISTTRASSSSRRFLSAITGVHGREIDSRGNPTVEVDVTTAQGT
>prePK-GFP *(Phaeodactylum)* (JGI 49002)
MMRSFLRHAQGRACAQHLRTIGTLRLNQMPVTGA
~~~

**C.**

~~~
>preTPI-GAPDH-GFP *(Phytophthora infestans)* (NCBI X64537)
MSFRQVFKTQARHMSSSSRKFFVGGNWKCNGSLGQAQELVGMLNTA
>prePGM-GFP *(Phytophthora infestans)* (PfGD Pi_011_55705_Feb05.seq)
MVLALRRPLAISSRVANRSLGMLRQQQKAMKHTHTLVLIRHGESEWNKKNLFTGWYDVQLSEKGNKEA
>prePK-GFP *(Achlya bisexualis)* (NCBI AAU81895)
MLARSLRSRAVRSFARGLSNKPSKNDAFSMT
>preTPI-GAPDH-GFP *(Saccharina latissima)* (NCBI ABU96661)
MFSAALSAAGAKAPSAARGFASSASRMSGRKFFVGGNWKCNGS
~~~

## *Phaeodactylum tricornutum* amino acid sequences used in GFP targeting experiments as seen in Supplementary Figure S4

~~~
>preTPI_50738 plastid (pre-sequence) (JGI 50738) plastid
MTGDSTSLLDLISPDRERPQRKEPSRWIAFSVFPFVRFIPEAFATRLPYSIVMKFLALSVAALISSATAFAPTFR
GSPASTTASTTSLAARKPFISGNWKLN
>preTPI_18228 plastid (pre-sequence) (JGI 18228) plastid
MKFLALSVAALISSATAFAPTFRGSPASTTASTTSLAARKPFISGNWKLN
>TPI_54738 cytosol (242 Amino acids) (JGI 54738) cytosol
MPRPDGSSTPAAEGERKYLVAGNWKCNGTLASNEELVKTFNEAGPIPSNVEVAICCPSLYLPQLLSSLRDDIQIG
AQDCGVNDKNGAFTGEIGAFQIKDIGCDWVIIGHSERRDGFEMPGETPDLCAKKTRVAIDAGLKVMFCIGEKKEQ
REDGTTMDVCASQLEPLAAVLTESDWSSIAIAYEPVWAIGTGLTATPEMAQETHASIRDWISQNVSADVAGKVRI
QYGGSMKGANAKDLLEQ
>Gapdh3_23 598 cytosol (full length) (JGI 23598) cytosol
MPVKCLVNGFGRIGRLCFRYAWDDPELEIVHVNDVCSCESAAYLVQYDSVHGTWSKSVVAAEDSQSFTVDGKLVT
FSQEKDFTKIDFASLGVDMVMECTGKFLTVKTLQPYFGMGVKQVVVSAPVKEDGALNVVLGCNHQKLTTDHTLVT
NASCTTNCLAPVVKVIQENFGIKHGCITTIHDVTGTQTLVDMPNTKKSDLRRARSGMTNLCPTSTGSATAIVEIY
PELKGKLNGLAVRVPLLNASLTDCVFEVNKEVTVEEVNAALKKASESGPLKGILGYETKPLVSTDYTNDTRSSII
DALSTQVIDKTMIKIYAWYDNEAGYSKRMAELCNIVAAMNITGQEPSFKYE
>prePGK_29157 plastid (pre-sequence) (JGI 29157) plastid
MKFVQAAIFALAASASTTAAFAPAKTFGVRSFAP
>PGK_51125 cytosol (193 Amino acids) (JGI 51125) cytosol
MASDMPKLAPGATRKRNVFDVIEALQKQSAKTILVRVDFNVPMNSDGKITDDSRIRGALPTIKAVVNAKCNAVLV
SHMGRPKLVQKAADDEETRQQRHELSLKPVADHLAKLLDQEVLFGDDCLHAQSTIRELPAEGGGVCLLENLRFYK
EEEKNGEDFRKTLASYADGYVNDAFGTSHRAHASVAGVPALLP
>PGM_43812 unclear (130 Amino acids) (JGI 43812) unclear localization,
cytosol plus ER or mitochondria
MGRRTTHRRLFPALALIFAELIMSTAYSLAWRTSAACWTTTTGTACSRSRIATTRKVRRSRPNPCNPWHPVAFSF
FGTSSRRCRSSGSLYGEIDADAEGPDSPSADDRSVPTPSTTSSLSRSETLPPIPP
>PGM_43253 mitochondria (112 Amino acids) (JGI 43253) mitochondria
MASITLNRSRFTMITAIGMSHPRSHGTPRSVLLLLLRQFSSKDWNSKGTDSASRSGPVLIKKTPRSAAAAKLRST
APSLNGSTTDSTTGAVKHHPAHHYINGGTPCDPAPPP
>PGM_26201 unclear (408 Amino acids) (JGI 26201) unclear, mitochondria or
ER
MLVPHPSGKAMRGLREEACRFLSSRSFGATLDATHARMGGNFVNSVQACNNGKRVCWHQRNRRTFSVVATQRNGI
GHRTTQGETEAVPRRHFTSLNQSTPFQLCFLRHGQSTWNRDNIFIGWTDTPLTDDGVLEARVAGKMLHKSGIRFD
EVHTSLLRRSIRTTNLALMELGQEYLPVHKHWRLNERCYGDLVGKNKKEVVMQHGADQVKRWRRSYDEPPPPMSD
DHPYHPARDPRYQNILDELPKSESLKNTVERSSLYWDEVLAPALREGKTLLVVGHENNLRSLLMRLEDIAPEDII
NLSLPRAVPLAYRLDENLKPLPREDGKLDEATGFLKGTWLGGDQAVSEILDRDHKQVYDTAITTNLEIGQDREKW
NNWMEFIMGKPSAKQKRIGGDKQNGFAGGAAIP
>PGM_42 8 57 plastid (175 Amino acids) (JGI 42857) plastid
MAMDAITMRKLTLTMAVLLIVSGCEALLVFLPRRSPFTVISTRSSTNSAGLLHLHSKANESDGLEGKWIKVSSAL
DEGVDAANEEKEGAFLSSDYNSMNGYNTDLNRYHTMLRERGTFVEALFGQRRSFVIAKRDGDENEDGWRDMRRQR
RPLWKHLLRLPISVAKNVLWKPPQP
>PGM_35164 mitochondria (351 Amino acids) (JGI 35164) mitochondria
MRIPCRRLHPQLSAKGTRRPFQYSSSNSIDDQHRSSHLDASPGRHIVVRHGQSVWNKGSNQLERFTGWTNVGLSE
NGQRQAVQAARKLHGYSIDCAYVSLLQRSQATLRLMLEELNDQGRRSEGYDDLTTDIPVISSWRLNERHYGALTG
QSKLQAEQLFGKAQLDLWRYSYKIPPPPMDPDTFSSWKHQAHCQMATYIHHRHNRSRVIEKGNSVWDSSRAVMPR
SEAFFDVLQRIVPLWKYGIAPRLARGETVLLVGHANSVKALLCLLDPHTVTPTSIGALKIPNTTPLVYQLIRDYP
GASTSVPASFPVLGDLRVVIPPSNSTRYPLSGTWLEDPPVARDAGTAVEEP
>PGM_51298 cytosol (131 Amino acids) (JGI 51298) cytosol. If a shorter
version, starting from the second Methionine is used, a localization at the
plastid as a blob like structure is the result (data not shown).
MCDESRQTATPMIHFEIFRFSDPLVRQDRQAPHLSLTSTVKILSDSNLHKLFIMMLRSLVLALSWTVASAFTHQS
TFWGRTAVTNSRILSLSPPTDASSSALCMKYMLVLVRHGESTWNKENRFTGWVDCP
>preENO_56468 cytosol (66 Amino acids) (JGI56468) cytosol. Start Methionine
from GFP was not included in the construct.
MLFKPSTLLALFAVAGTTLAFAPRSTTTPLTSTTRGSASSSVTTLAMSGITGVLAREILDSRGNPV
>ENO_56468 plastid (443 Amino acids) (JGI56468) plastid
MLFKPSTLLALFAVAGTTLAFAPRSTTTPLTSTTRGSASSSVTTLAMSGITGVLAREILDSRGNPTVEVEVTTAD
GVFRASVPSGASTDAYEAVELRDGGDRYMGKGVLQAVQNVNDILGPAVMGMDPVGQGSVDDVMLELDGTPNKANL
GANAILGVSLAVAKAGAAAKKVPLYRHFADLAGNNLDTYTMPVPCFNVINGGSHAGNKLAFQEYFVIPTGAKSFA
EAMQIGCEVYHTLGKIIKAKFGGDATLIGDEGGFAPPCDNREGCELIMEAISKAGYDGKCKIGLDVAASEFKVKG
KDEYDLDFKYDGDIVSGEELGNLYQSLAADFPIVTIEDPFDEDDWENWSKFTTKNGATFQVVGDDLTVTNIEKIE
RAIDEKACTCLLLKVNQIGSISESIAAVTKAKKAGWGVMTSHRSGETEDTYIADLAVGLCTGQIKTGA
>PK_49098 cytosol (507 Amino acids) (JGI: 49098) cytosol
MTASQTKITASGPELRGANITLDTIMKKTDVSTRQTKIVCTLGPACWEVEQLESLIDAGLSIARFNFSHGDHEGH
KACLDRLRQAADHKKKHVAVMLDTKGPEIRSGFFADGAKKISLVKGETIVLTSDYSFKGDKHKLACSYPVLAKSV
TPGQQILVADGSLVLTVLSCDEAAGEVSCRIENNAGIGERKNMNLPGVIVDLPTLTDKDIDDIQNWGIVNDIDFI
AASFVRKASDVHKIREVLGEKGKGIKIICKIENQEGMDNYDEILEATDAIMVARGDLGMEIPPEKVFLAQKMMIR
QANIAGKPVVTATQMLESMITNPRPTRAECSDVANAVLDGTDCVMLSGETANGEYPTAAVTIMSETCCEAEGAQN
TNMLYQAVRNSTLSQYGILSTSESIASSAAKTAIDVGAKAIIVCSESGMTATQVAKFRPGRPIHVLTHDVRVARQ
CSGYLRGASVEVISSMDQMDPAIDAYIERCKANGKAVAGDAFVVVTGTVAQRGVTNA
>PK_56445 cytosol (538 Amino acids) (JGI 56445) cytosol
MSLSQSSDVPILAGGFITLDTVKHPTNTINRRTKIVCTIGPACWNVDQLEILIESGMNVARFNFSHGDHAGHGAV
LERVRQAAQNKGRNIAILLDTKGPEIRTGFFANGASKIELVKGETIVLTSDYKFKGDQHKLACSYPALAQSVTQG
QQILVADGSLVLTVLQTDEAAGEVSCRIDNNASMGERKNMNLPGVKVDLPTFTEKDVDDIVNFGIKHKVDFIAAS
FVRKQSDVANLRQLLAENGGQQIKICCKIENQEGLENYDEILQATDSIMVARGDLGMEIPPAKVFLAQKMMIREA
NIAGKPVITATQMLESMINNPRPTRAECSDVANAVLDGTDCVMLSGETANGPYFEEAVKVMARTCCEAENSRNYN
SLYSAVRSSVMAKYGSVPPEESLASSAVKTAIDVNARLILVLSESGMTAGYVSKFRPERAIVCLTPSDAVARQTG
GILKGVHSYVVDNLDNTEELIAETGVEAVKAGIASVGDLMVVVSGTLYGIGKNNQVRVSVIEAPEGTVKETPAAM
KRLVSFVYAADEI
>PK_45997 cytosol (533 Amino acids) (JGI 45997) cytosol
MLSSTSTIPKLDGEVVTLSIIKKPTETKKRRTKIICTLGPACWSEEGLGQLMDAGMNVARFNFSHGDHEGHGKVL
ERLRKVAKEKKRNIAVLLDTKGPEIRTGFFADGIDKINLSKGDTIVLTTDYDFKGDSKRLACSYPTLAKSVTQGQ
AILIADGSLVLTVLSIDTANNEVQCRVENNASIGERKNMNLPGVVVDLPTFTERDVNDIVNFGIKSKVDFIAASF
VRKGSDVTNLRKLLADNGGPQIKIICKIENQEGLENYGDILEHTDAIMVARGDLGMEIPSSKVFLAQKYMIREAN
VAGKPVVTATQMLESMVTNPRPTRAECSDVANAVYDGTDAVMLSGETANGPHFEKAVLVMARTCCEAESSRNYNL
LFQSVRNSIVIARGGLSTGESMASSAVKSALDIEAKLIVVMSETGKMGNYVAKFRPGLSVLCMTPNETAARQASG
LLLGMHTVVVDSLEKSEELVEELNYELVQSNFLKPGDKMVVIAGRMAGMKEQLRIVTLDEGKSYGHIVSGTSFFF
ERTRLLDF
>PK_56172 mitochondria (86 Amino acids) (JGI: 56172) mitochondria
MFRRAVLSLSTRAIRTPVPCSVARGDASQVRSLAQTTFYLPDPADRSQDVHNRGNLQLSKIVATIGPTSEQEEPL
RLVTDAGMRIM
~~~

## Supplementary Tables and Figures

**Supplementary Table 1.** *Blastocystis* glycolytic enzymes are protected by a membrane. Control: Mitochondrial fractions incubated without proteolytic enzymes. Protease: Mitochondria incubated in 225 mM sucrose buffer at 25 ^o^C containing 500 U bovine pancreas trypsin, 10 U papaya latex papain and 250 U porcine pepsin for 15 minutes. Protease + Triton: Mitochondrial fractions containing proteolytic enzymes and 1% Triton X-100 incubated for 15 min at 25 ^o^C. Samples were centrifuged (14,000 g) for 2 min and resuspended in fresh sucrose buffer without proteolytic enzymes prior to assay.

|  | control | protease | protease + Triton |
| --- | --- | --- | --- |
| glyceraldehyde phosphate dehydrogenase | 71 ± 14 | 61 ± 9.1 | 15 ± 4.8 |
| phosphoglycerate kinase | 36 ± 8.5 | 33 ± 4.5 | 6.3 ± 2.2 |
| phosphoglycerate mutase | 4.0 ± 1.2 | 3.1 ± 0.5 | 0.2 ± 0.1 |
| enolase | 1.8 ± 0.9 | 1.4 ± 1.0 | 0.3 ± 0.1 |
| phosphoenolpyruvate synthase | 28 ± 5.1 | 25 ± 4.9 | 7.6 ± 2.0 |

(1 μmol substrate to product per minute)

**Figure S1.**
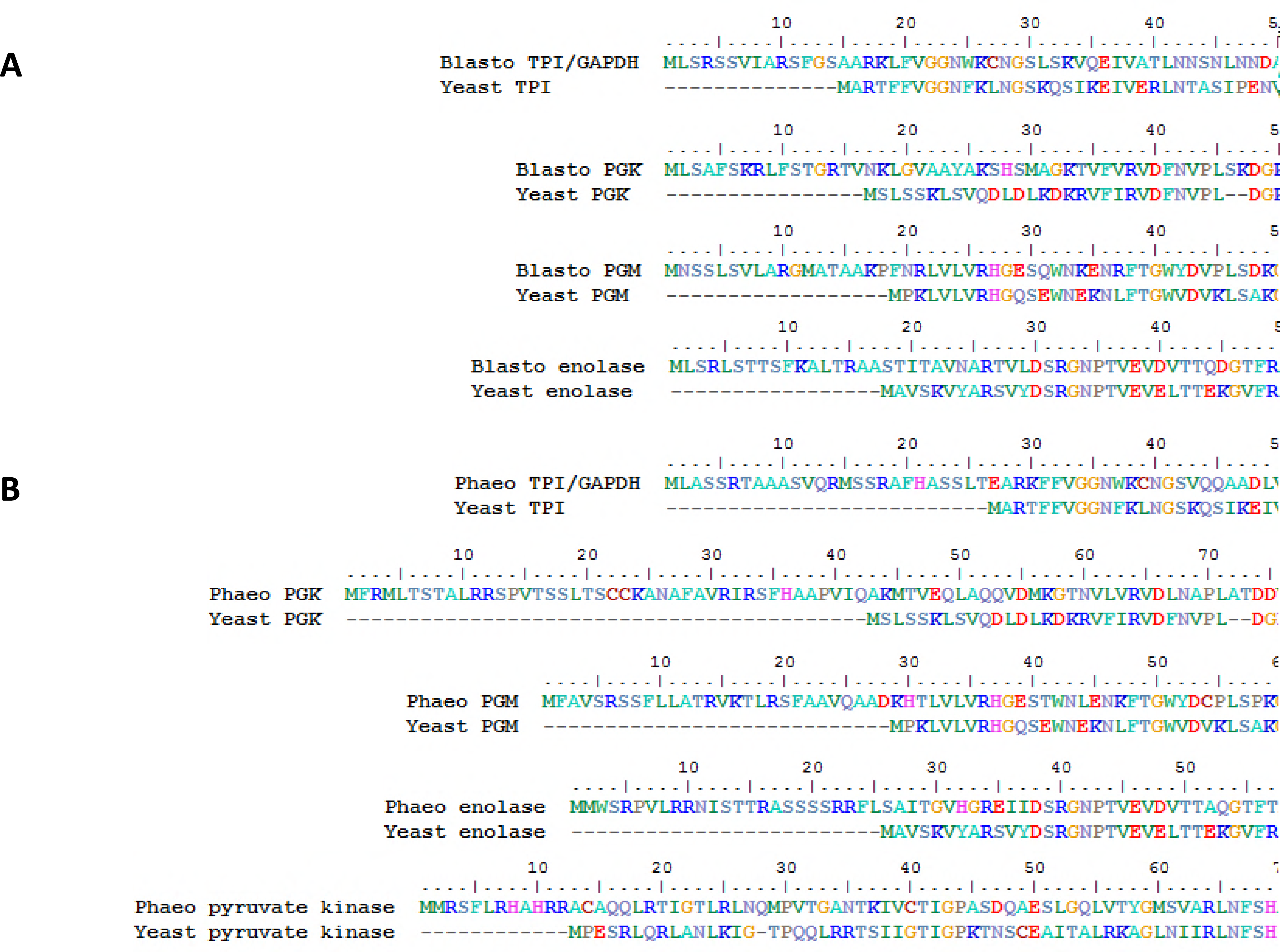
Stramenopile glycolytic C3 enzymes contain amino-terminal targeting signals. A. Comparison of *Blastocystis* amino-terminal sequences for TPI-GAPDH, PGK, PMG, and enolase with homologs from yeast showing the mitochondrial-like targeting signals. B. *Phaeodactylum tricornutum* glycolytic C3 enzyme amino-termini of TPI-GAPDH, PGK, PMG, enolase, and pyruvate kinase compared to yeast homologs demonstrate mitochondrial-like targeting signals.

**Figure S2.**
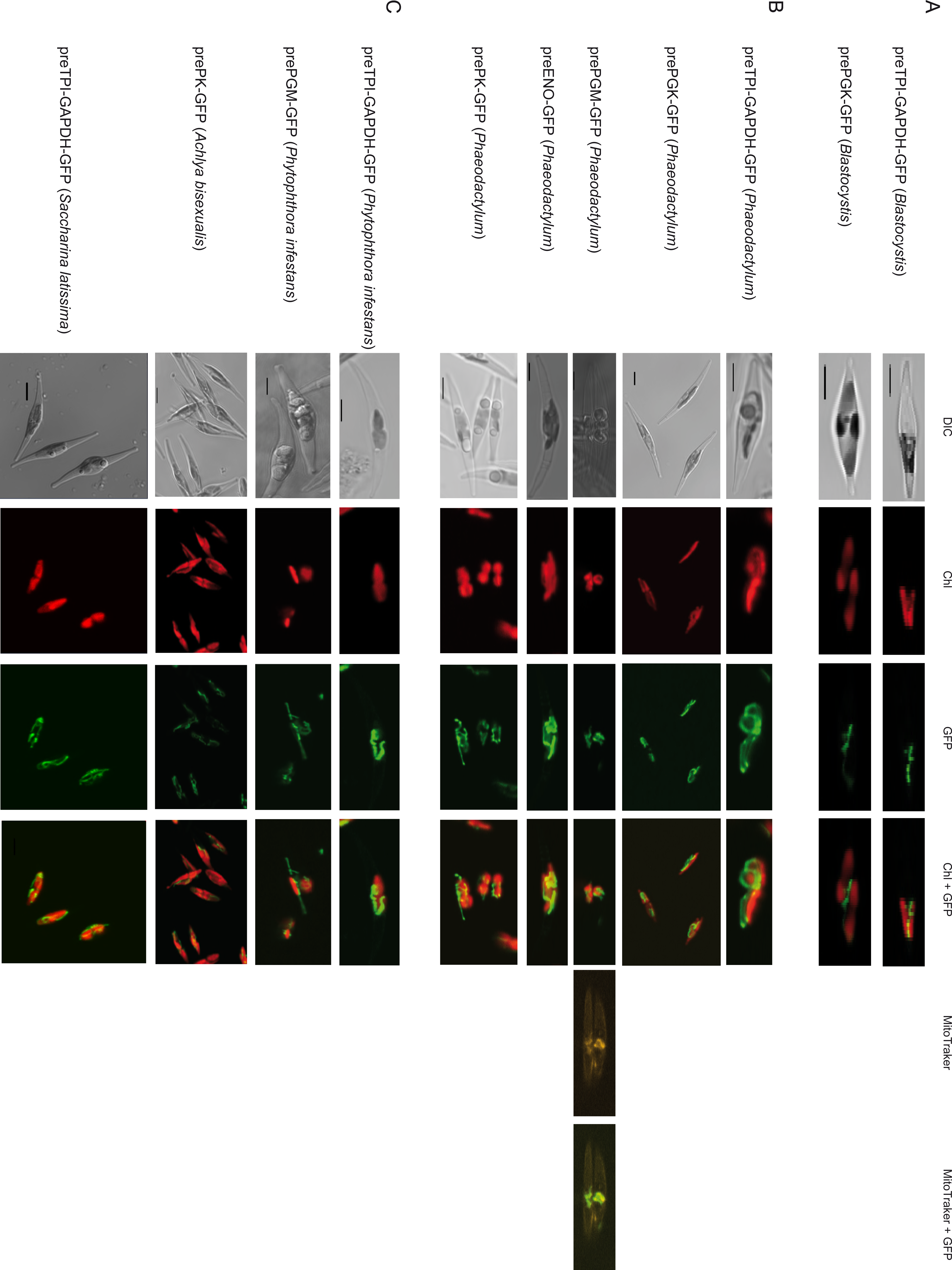
Stramenopile glycolytic enzyme amino-terminal mitochondrial-like targeting signals are sufficient to target GFP to mitochondria in the diatom *Phaeodactylum tricornutum.* A. The *Blastocystis* glycolytic enzymes TPI-GAPDH and PGK contain amino-terminal targeting signals that can target GFP to *P. tricornutum* mitochondria. B. Amino-terminal extensions on TPI-GAPDH, PGK, phosphoglycerate mutase (PGM), enolase and pyruvate kinase (PK) from the diatom *P. tricornutum* were cloned in front of GFP and constructs used to transform *P. tricornutum.* C. Amino terminal extensions on TPI-GAPDH and PGM from *Phytophthora infestans*, PK from *Achlya bisexualis* and TPI-GAPDH from *Saccharina latissima* were used as above to test for functionality of targeting information in *P. tricornutum.* DIC, Differential interference contrast microscopy. Chl, Chlorophyll *a* autofluorescence. GFP, Green fluorescent protein. Chl+GFP, Merged imaged showing the discrete (mitochondrial) localization of GFP. MitoTraker, MitoTraker Orange stain. MitoTraker+GFP, Merged image show considerable overlap of MitoTraker stain and GFP fluorescence. For the corresponding amino acid sequences used for GFP targeting, see Supplementary File 1. Scale bar 5 μm.

**Figure S3.**
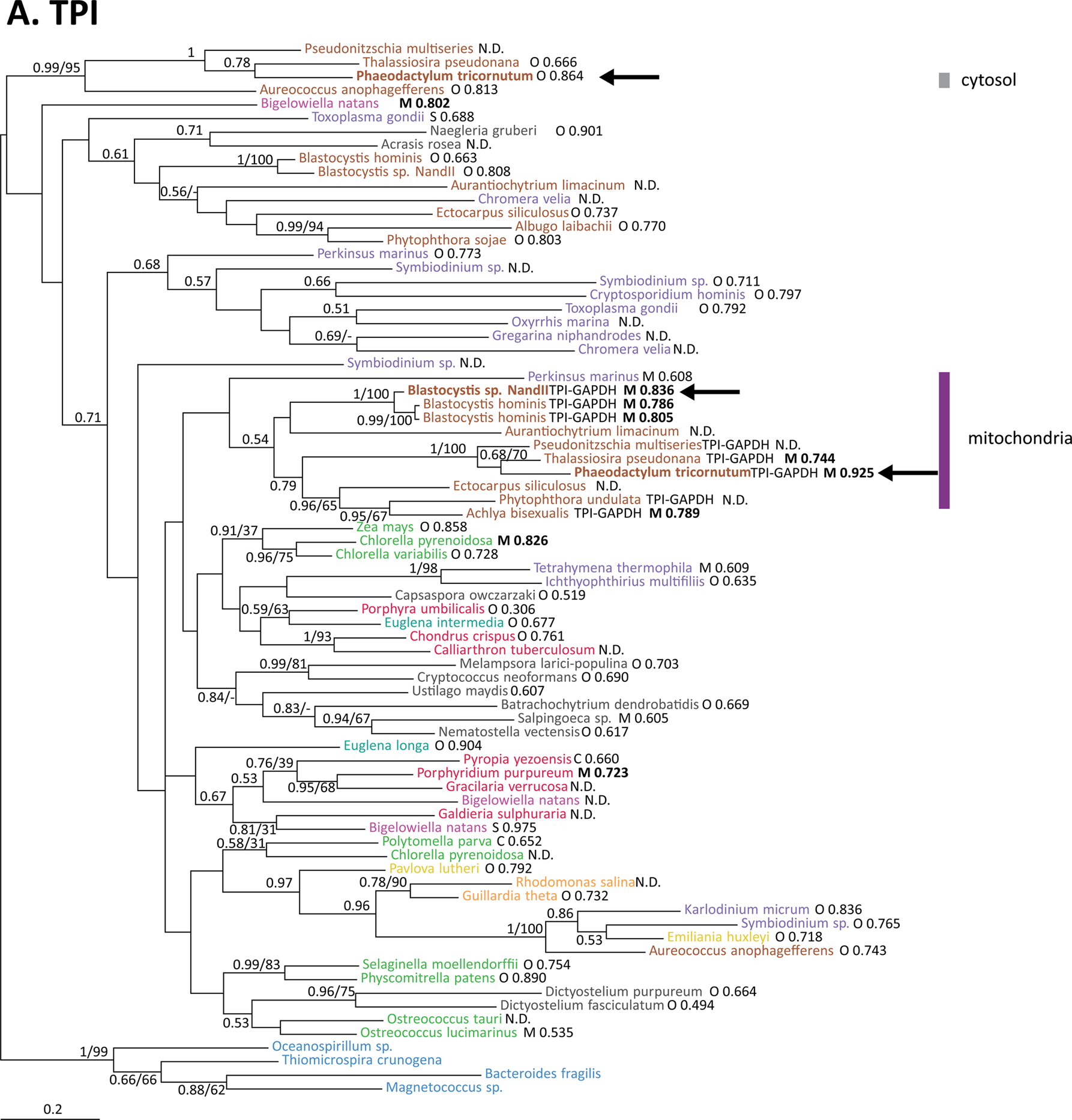
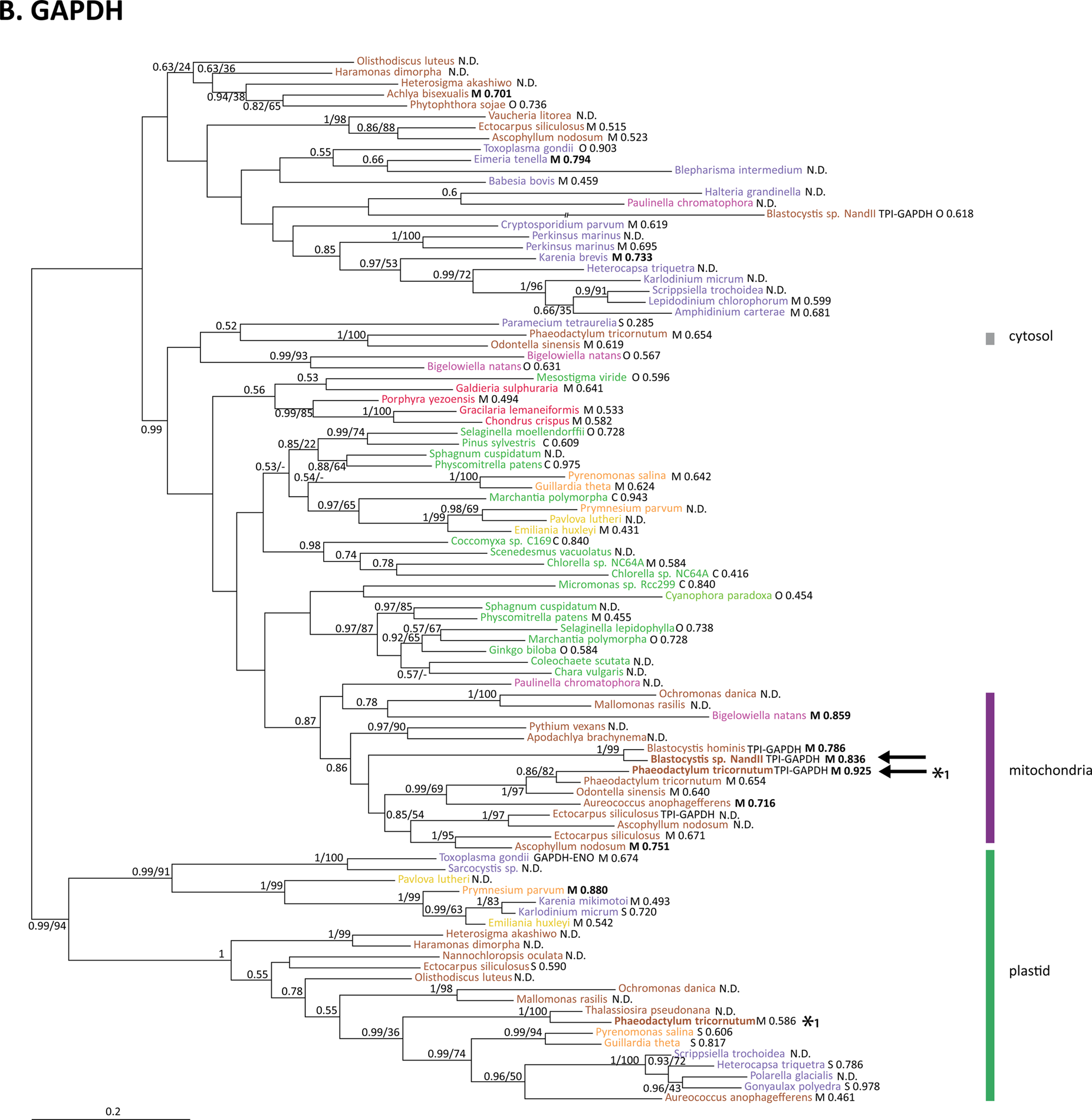
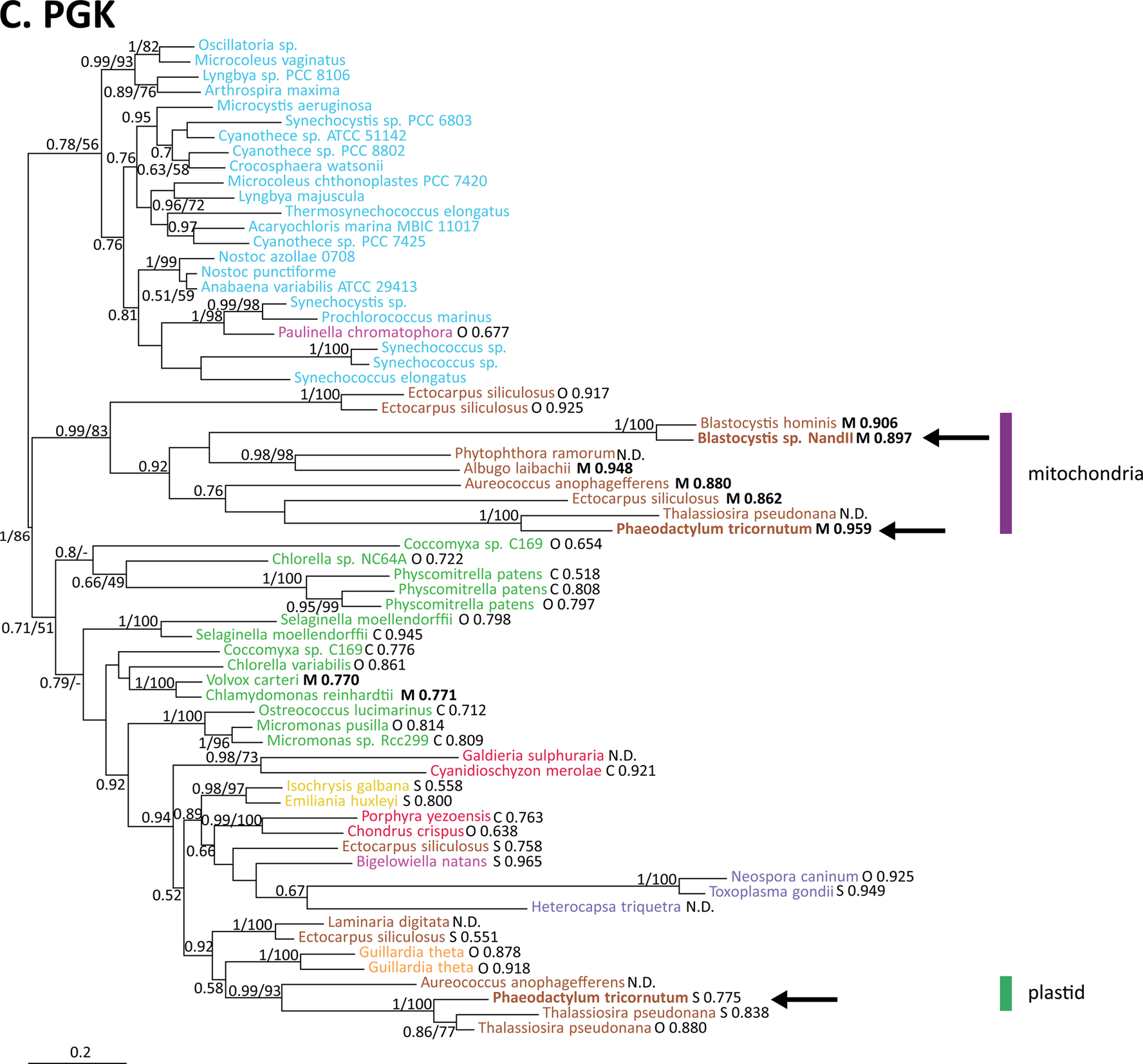
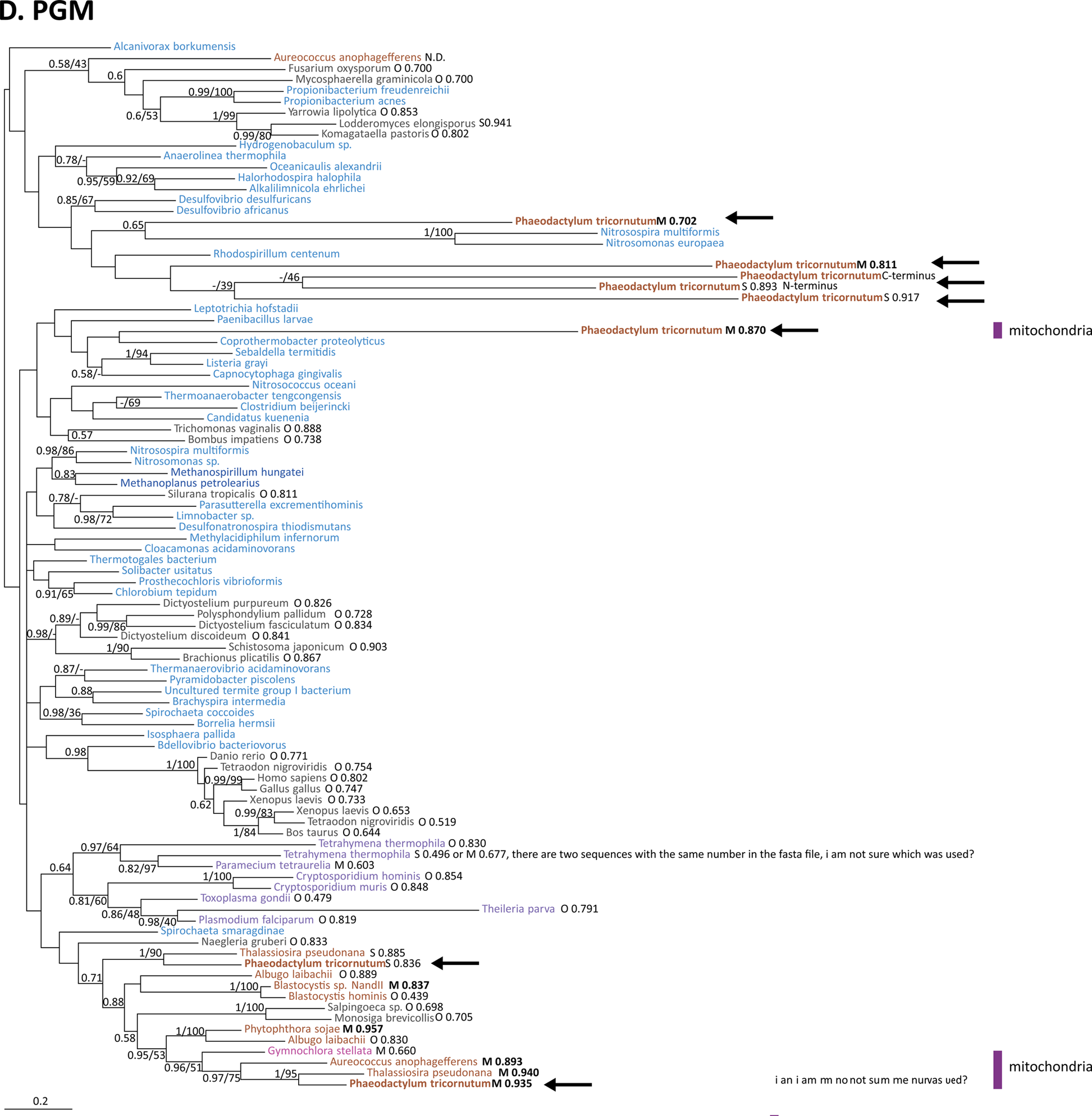
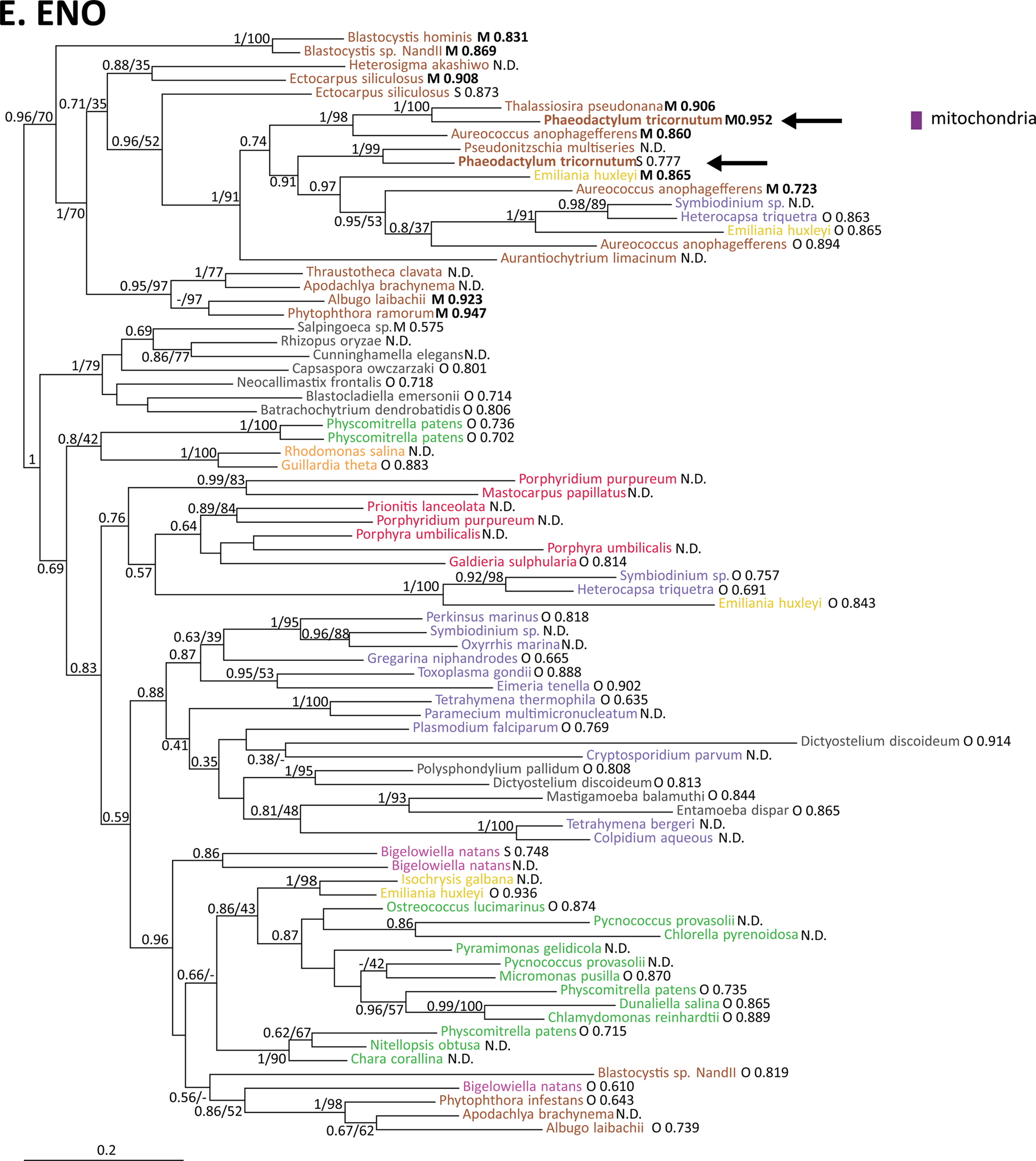
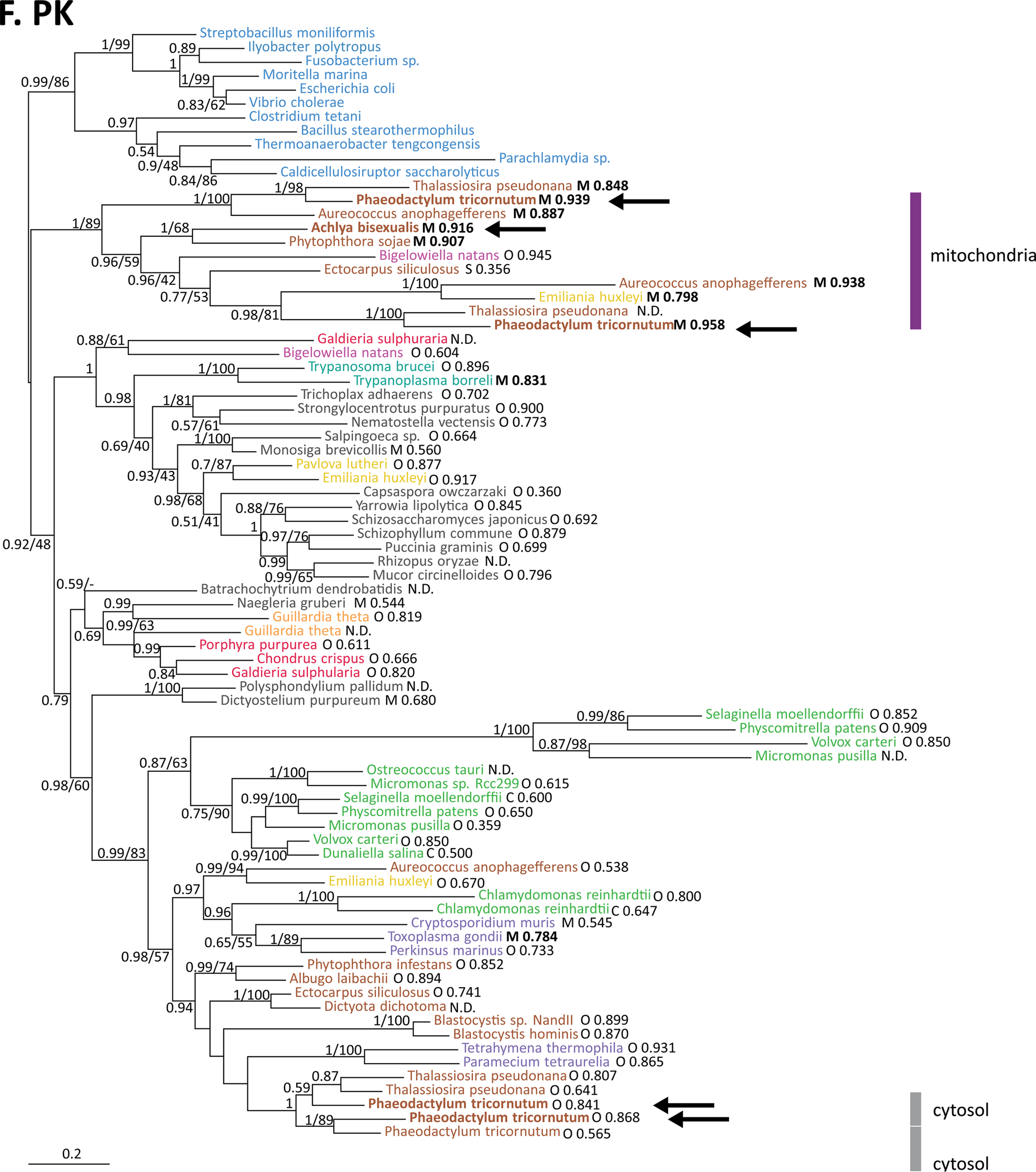
Phylogenetic analysis of glycolytic enzymes of the pay-off phase. A. Triosephosphate isomerase (TPI), 77 sequences and 167 amino acid positions were used to calculate the tree. Bacterial sequences were used as outgroup. B. Glyceraldehyde-3-phosphate dehydrogenase (GAPDH), only GAPDH from the C-type are illustrated in the tree. 96 sequences and 266 amino acid positions were used to calculate the tree. One branch caused long branch attraction (LBA) artifact and therefore is shortened in the figure. C. Phosphoglycerate kinase (PGK), 66 sequences and 360 amino acid positions were used to calculate the tree. D. Phosphoglycerate mutase (PGM), 96 sequences and 199 amino acid positions were used to calculate the tree. E. Enolase (ENO), 80 sequences and 345 amino acid positions were used to calculate the tree. F. Pyruvate kinase (PK), 80 sequences and 287 amino acid positions were used to calculate the tree. Values at nodes: posterior probabilities (P.P. >0.5) / rapid bootstrap values (BS>30%). Species name in bold = localization experimental proof (arrow = in this study, star Liaud *et. al* (2000)). TargetP analysis: M = mTP = mitochondria (bold indicates scores > 0.700), O = other, SP = signal peptide, C = cTP = chloroplast only for Viridiplantae, Rhodophyta and Glaucocystophyceae plant results was taken if non-plant results differ. N.D.: sequences not analyzed; not complete at N-terminus or start methionine is missing (proof by an alignment). Colour code: Viridiplantae, Stramenopiles, Alveolata, Rhizaria, Rhodophyta, Cryptophyta, Haptophyceae, Bacteria, other Eukaryota, Archaea, Cyanobacteria, Euglenozoa.

**Figure S4.**
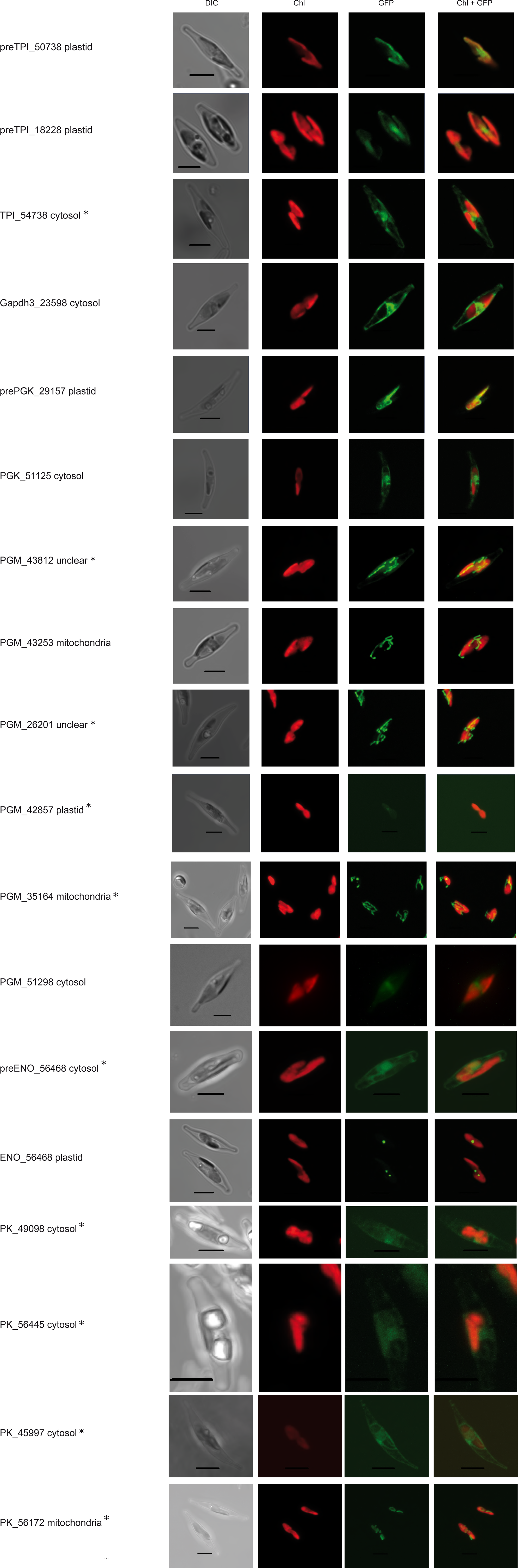
*Phaeodactylum tricornutum* contains, similar to some other stramenopiles, multiple isoforms for the C3 part of glycolysis. The localization for all isoforms was tested via GFP-fusion constructs. A “pre” suffix means that the predicted targeting signal was used; if the suffix is missing the full length of the respective sequence was fused to GFP. The number is the JGI Protein ID and the result of each localization is mentioned. For the corresponding amino acid sequences used for GFP targeting see Supplementary File 2. A star (*) marks images were a maximum intensity projection from a Z-Stack was used. Unclear indicates localization not possible to identify. TPI = Triosephosphate isomerase, GAPDH = Glyceraldehyde-3-phosphate dehydrogenase, PGK = Phosphoglycerate kinase, PGM = Phosphoglycerate mutase, ENO = Enolase, PK = Pyruvate kinase. DIC, Differential interference contrast microscopy. Chl, Chlorophyll *a* autofluorescence. GFP, Green fluorescent protein. Chl+GFP, Merged imaged showing the discrete localization of GFP compared to Chlorophyll autoflourescence. For the corresponding amino acid sequences used for GFP targeting, see Supplementary File 2. Scale bar 5 μm.

